# Regulation of the MDM2-p53 Nexus by a Nuclear Phosphoinositide and Small Heat Shock Protein Complex

**DOI:** 10.1101/2025.04.11.648454

**Authors:** Jeong Hyo Lee, Mo Chen, Tianmu Wen, Richard A. Anderson, Vincent L. Cryns

## Abstract

The tumor suppressor p53 maintains genome stability in the setting of cellular stress and is frequently mutated in cancer. The stability of p53 is regulated by its interaction with the oncoprotein MDM2, a ubiquitin E3 ligase. Recently, nuclear phosphoinositides were reported to bind and stabilize p53. Here, we report that genotoxic stress induces the type I phosphatidylinositol phosphate kinase (PIPKIα) and its product phosphatidylinositol 4,5-bisphosphate (PIP_2_) to bind and regulate the stability and function of MDM2. Following genotoxic stress, nuclear PIPKIα binds to MDM2 to generate a complex of MDM2 and PIP_2_. PIP_2_ binding to MDM2 differentially regulates the recruitment of the small heat shock proteins (sHSPs) αB-crystallin (αBC) and HSP27 to the MDM2-PIP_2_ complex, acting as an on-off switch that regulates MDM2 stability, downstream targets, ubiquitination activity, and interaction with p53. Our results demonstrate an unexpected role for nuclear phosphoinositides conferring specificity to the MDM2-PIP_2_-sHSPs association. Notably, the differential engagement of αBC and HSP27 reveals that sHSPs are not merely passive chaperones but play active, selective roles in fine-tuning MDM2 function and MDM2-p53 nexus. These findings provide a novel therapeutic strategy for targeting this pathway in cancer.

## Introduction

Phosphatidylinositol-4,5-bisphosphate (PIP_2_) is the most abundant of the seven phosphoinositide lipid second messengers and plays a crucial role in regulating various cellular functions, including membrane trafficking, ion channel activity, and the actin cytoskeleton (1). PIP_2_ directly interacts with PIP_2_ effector proteins, controlling their activity and/or localization (2, 3). PIP_2_ is predominantly synthesized by the phosphorylation of phosphatidylinositol-4-phosphate (PI4P) and phosphatidylinositol-5-phosphate (PI5P) by type I (PI4P 5-kinases) and type II (PI5P 4-kinases) phosphatidylinositol phosphate kinases (PIPKs), respectively. The type I and type II kinases have α, β, and γ isoforms. PIPKIα, PIPKIβ, PIPKIγ, PIPKIIα and PIPKIIβ have reported nuclear activities (4). PIP_2_ is present in the nucleus as well as most membrane structures, including the plasma membrane and membranes of the endosome, Golgi, and endoplasmic reticulum. In the nucleus, PIP_2_ localizes to regions distinct from the nuclear envelope in non-membranous nuclear speckles (5). Although several nuclear PIP_2_ effectors have been identified (2, 3, 5), the function of PIP_2_ in the nucleus is poorly understood.

One such nuclear PIP_2_ effector recently reported is the p53 tumor suppressor protein (wild-type and mutant forms)^3^. Wild-type p53 defends the genome from various cellular stressors and is the most frequently mutated gene in human tumors (6, 7). p53 mutations result in the loss of wild-type p53 activities and gain of function, which promote cell survival and stress resistance to drive cancer initiation/progression (8–11). Remarkably, PIPKIα and its product PIP_2_ interact with the C-terminal domain (CTD) of stress-activated wild-type p53 and mutant p53, resulting in the formation of highly stable p53-PIP_2_ complexes in the nucleus that are resistant to harsh denaturing conditions. These complexes then recruit the small heat shock proteins (sHSPs) αB-Crystallin (αBC) and HSP27, which bind and stabilize p53 (3). Moreover, the nuclear PI3K inositol polyphosphate multikinase (IPMK) and the 3-phosphatase phosphatase and tensin homolog (PTEN) bind p53 to catalyze the synthesis of p53-PIP_2_ to p53-phosphatidylinositol-3,4,5-triphosphate (PIP_3_) and the reverse reaction, respectively (12). p53-PIP_3_ recruits and activates Akt in the nucleus by a PIP_3_-dependent mechanism, providing a direct functional link between p53 and Akt in the nucleus. These findings indicate that phosphoinositides and PIP kinases dynamically and spatially regulate p53 stability and function in the nucleus.

Given the central role of the E3 ubiquitin-protein ligase, MDM2, in regulating p53 stability by ubiquitin-dependent proteasomal degradation of p53 (13, 14), we postulated that phosphoinositides and sHSPs might regulate the MDM2-p53 interaction by associating with p53 and/or MDM2. Indeed, both HSP90 and HSP70 bind mutant p53 and negatively regulate p53 degradation by MDM2 (15–17). Here, we report that MDM2, like p53, directly binds PIPKIα and its product PIP_2_, leading to the recruitment of sHSPs to MDM2, which regulates its stability, E3 ligase activity, and interaction with p53. Taken together, both MDM2 and p53 are regulated by phosphoinositides, thereby providing exquisite temporal and spatial regulation of MDM2-p53 interaction in the nucleus.

## Results

### PIP_2_ associates with MDM2 in the nucleus in response to stress

As MDM2 has clusters of positively charged amino acids consistent with one or more putative phosphoinositide interaction motifs (2, 18, 19), we utilized a liposome sedimentation assay to investigate MDM2 binding to multiple phosphatidylinositol phosphate (PIP_n_) isomers. Recombinant MDM2 protein was incubated with PIP_n_-beads and MDM2-PIP_n_ liposomes, which were sedimented and analyzed by immunoblotting for MDM2. Although phosphatidylinositol (PI) did not bind MDM2, PI4P, PIP_2,_ and PIP_3_ bound to MDM2, with PIP_2_ exhibiting the most robust interaction (Fig. 1*A*). The specificity of the MDM2 antibody used throughout this study was validated by MDM2 KD in multiple cell lines (Fig. S1*A*) consistent with prior reports (20). Microscale thermophoresis (MST) confirmed direct binding of PIP_2_ to MDM2 (K_d_ 352 ± 3 nM, Table 1*A*). Consistent with their binding *in vitro*, cisplatin treatment increased nuclear levels of MDM2 and PIP_2_ and their nuclear co-localization as determined by immunofluorescence (IF) staining (Fig. 1*B*). Additionally, genotoxic stress increased the amount of PIP_2_ that co-immunoprecipitated with MDM2 across a panel of mutant p53 or p53-null cancer cell lines, while this interaction was decreased by cisplatin in a p53 wild-type cell line (Fig. 1*C* and Fig. S1*B*). Proximity ligation assay (PLA), which detects two closely located epitopes within 40 nm, revealed that the majority of the MDM2-PIP_2_ and p53-MDM2 foci were localized in the nucleus, with both signals increasing in response to cisplatin (Fig. 1*D* and Fig. S1*C*). Furthermore, we confirmed the interaction of MDM2 and PIP_2_ by metabolic labeling cells with [^3^H]*my*o-inositol, which resulted in the incorporation of [^3^H]*my*o-inositol into MDM2. Liquid scintillation counting (LSC) demonstrated that [^3^H]*my*o-inositol was primarily detected in 75∼100 kDa gel slices, corresponding to the molecular weight of MDM2, in the [^3^H]*my*o-inositol-treated group (Fig. 1*E*). Notably, this incorporation of [^3^H]*my*o-inositol into MDM2 was not disrupted by harsh denaturing conditions and was enhanced by cisplatin treatment. Collectively, these data indicate that PIP_2_ directly binds to MDM2 with high affinity *in vitro* and stably associates with MDM2 in the nucleus of cells in response to stress.

**Fig. 1:**
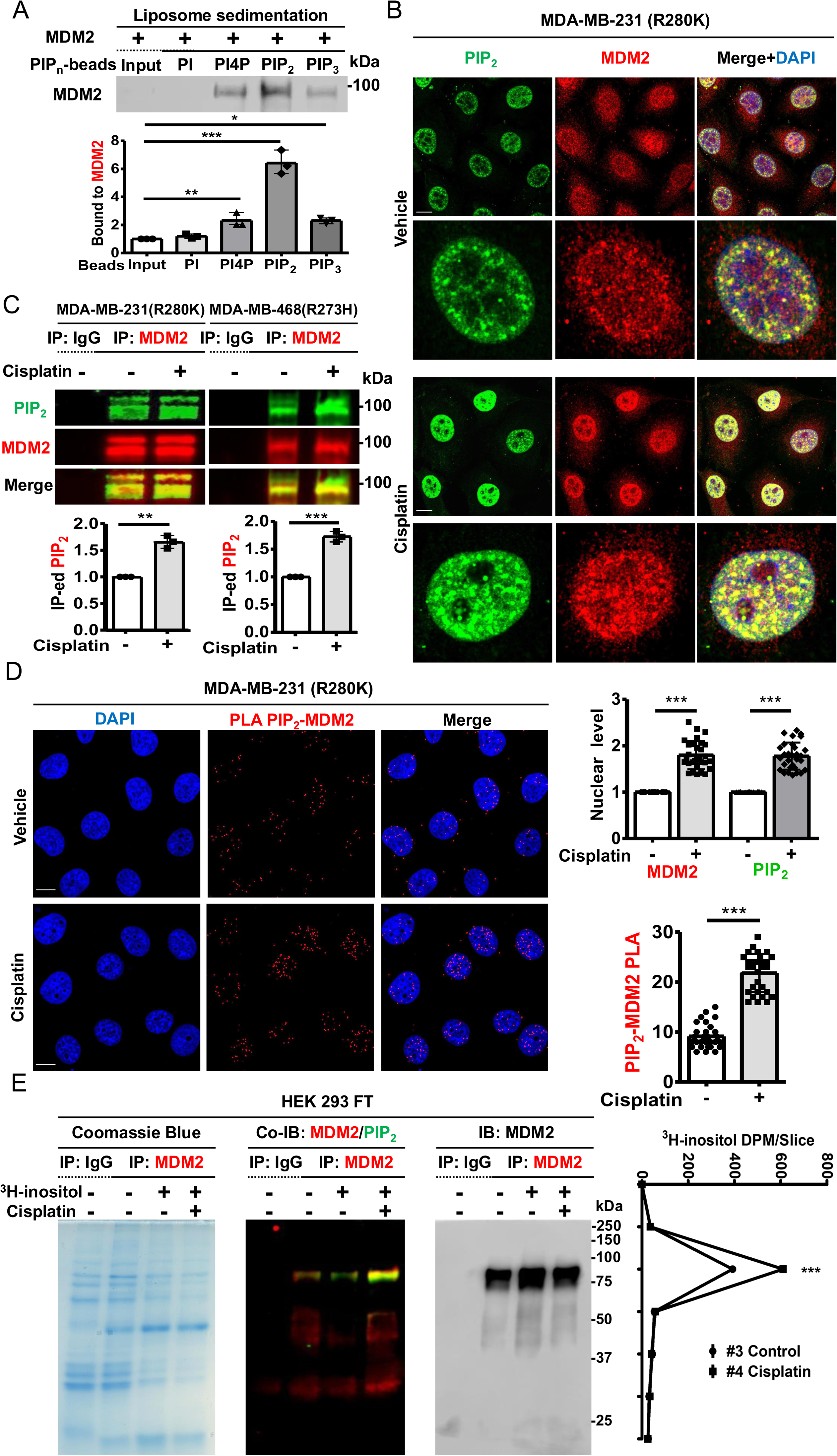
PIP_2_ associates with MDM2 in the nucleus in response to stress. ***A***, MDM2 recombinant protein (0.1Lµg) was incubated with 1.0LnM liposomes containing the indicated phosphoinositides. Liposomes were sedimented, and the associated MDM2 was analyzed by immunoblotting (IB). The MDM2 IB intensity was quantified, and the graph is shown as meanL± standard deviation(s.d.) of *n*L=L4 independent experiments ***B***, Confocal images of immunofluorescence (IF) staining against PIP_2_ and MDM2 in MDA-MB-231 cells treated with vehicle or 30LµM cisplatin (Cis) for 24Lh. MDM2 and PIP_2_ antibodies were used for the immunostaining study: scale bars, 5 µm. ***C***, MDA-MB-231, and MDA-MB-468 cells were treated with 30LµM cisplatin or vehicle for 24Lh and then processed for immunoprecipitation (IP) of MDM2 and fluorescence immunoblot (IB). Fluorescence IP−IB detects stress-induced PIP_2_ association with endogenous MDM2. The PIP_2_ IB intensity was quantified, and the graph is shown as meanL±Ls.d. of *n*L=L3 independent experiments. ***D***, Proximity ligation assay (PLA) of MDM2-PIP_2_ in MDA-MB-231 cells treated with vehicle or 30LµM cisplatin for 24Lh. The nuclear PLA foci of MDM2-PIP_2_ were quantified. *n*L=L30 cells pooled from 3 independent experiments, 10 cells per experiment ***E***, HEK293FT cells transiently transfected with MDM2 were treated with vehicle or 25 µCi/ml ^3^H-myo-inositol to metabolic labeling. Cells were treated with vehicle or 30LµM cisplatin for 24Lh, then processed for Coomassie blue staining and IP of MDM2 and fluorescence IB. The gel was sliced by molecular weight corresponding to the adjacent graph and analyzed by liquid scintillation counting (LSC). *n*L=L3 slices pooled from 3 independent experiments, 6 slices per experiment.

**Table 1:**
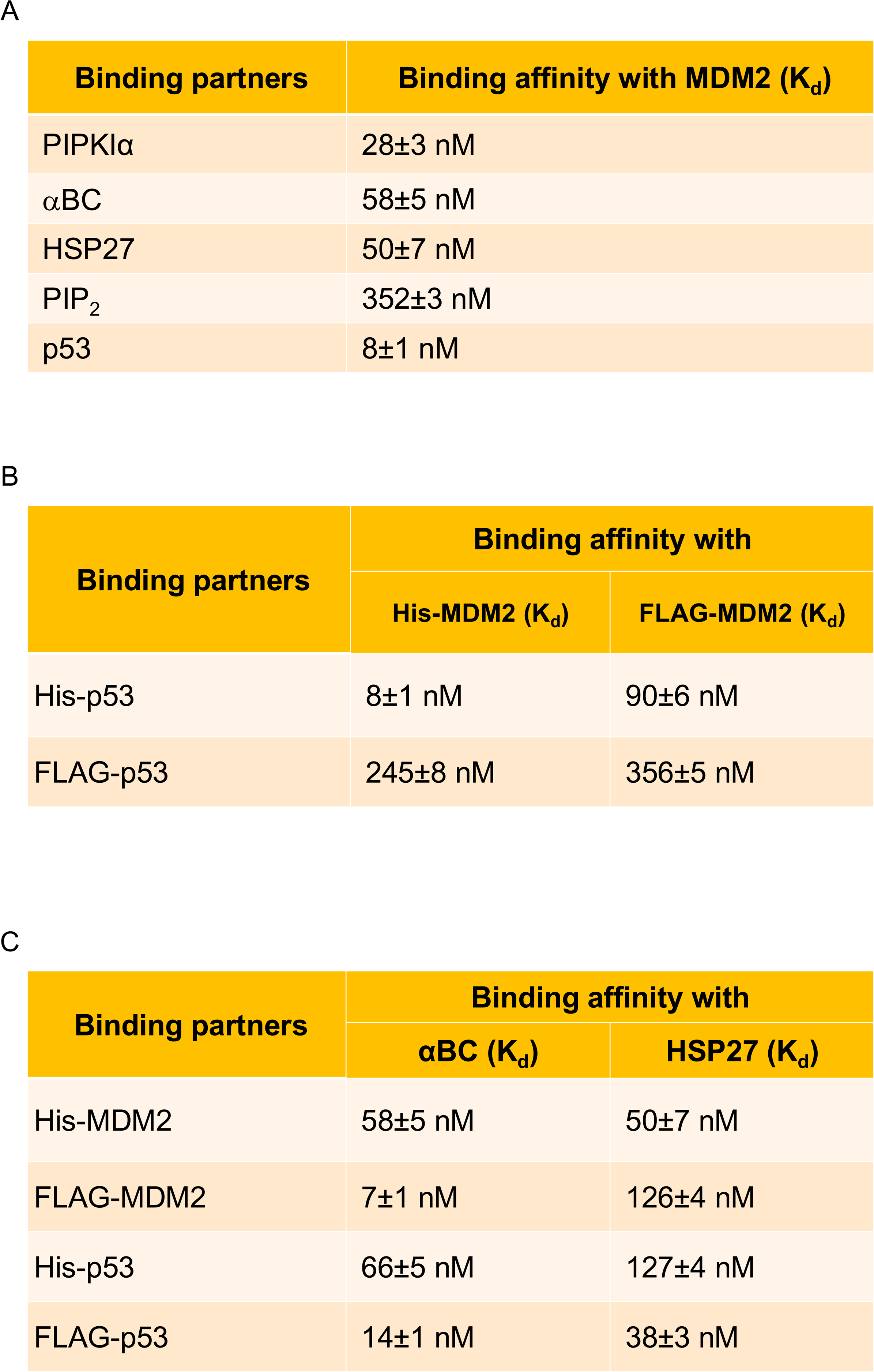
MST Analysis of the binding affinity of MDM2 to PIPKIα, αBC, HSP27, PIP_2_ and p53. *A*, Interaction of fluorescence-labeled MDM2 and non-labeled PIPKIα, αBC, HSP27, PIP_2_ and p53 was evaluated using microscale thermophoresis (MST). MST traces of a constant concentration of fluorescence-labeled MDM2 (target, 5 nM) incubated with increasing concentrations to 500 nM of PIPKIα, αBC, HSP27, PIP_2_ and p53 (ligand) were measured and used for calculating the binding affinity. MST revealed direct binding of MDM2 to PIPKIα, αBC, HSP27, PIP_2_, and p53 at indicated K_d_ values. MST was performed using a Monolith NT.115 Pico, and the binding affinity was auto-generate using Control v1.6 software. nL=L3 independent experiments. ***B***, Binding affinity of fluorescence-labeled bacteria-expressed His_MDM2 and mammalian-expressed FLAG-MDM2 (target, 5 nM) with gradually increasing amounts to 500 nM of non-labeled His-p53 and FLAG-p53 (ligand) was calculated using MST assay. MST was performed using a Monolith NT.115 Pico, and the binding affinity was auto-generated. Control v1.6 software. nL=L3 independent experiments. ***C*,** Direct binding of fluorescence-labeled αBC and HSP27 (target, 5 nM) with increasing amounts to 500 nM of non-labeled His-MDM2, FLAG-MDM2, His-p53 and FLAG-p53 (ligand) was analyzed using MST as in *A*.

| Binding partners | Binding affinity with MDM2 (K<sub>d</sub>) |
| --- | --- |
| PIPKIα | 28±3 nM |
| αBC | 58±5 nM |
| HSP27 | 50±7 nM |
| PIP<sub>2</sub> | 352±3 nM |
| p53 | 8±1 nM |

B
| Binding partners | Binding affinity with |  |
| --- | --- | --- |
|  | His-MDM2 (K<sub>d</sub>) | FLAG-MDM2 (K<sub>d</sub>) |
| His-p53 | 8±1 nM | 90±6 nM |
| FLAG-p53 | 245±8 nM | 356±5 nM |

C
| Binding partners | Binding affinity with |  |
| --- | --- | --- |
|  | αBC (K<sub>d</sub>) | HSP27 (K<sub>d</sub>) |
| His-MDM2 | 58±5 nM | 50±7 nM |
| FLAG-MDM2 | 7±1 nM | 126±4 nM |
| His-p53 | 66±5 nM | 127±4 nM |
| FLAG-p53 | 14±1 nM | 38±3 nM |

### PIPKI**α** interacts with MDM2 and regulates PIP_2_ association

As PIPKIα binds p53 and transfers its product PIP_2_ to p53, (2, 3, 12) we examined its potential functional role in transferring PIP_2_ to MDM2. Consistent with this idea, PIPKIα co-IPed with MDM2 in a panel of wild-type p53, mutant p53, and p53-null cancer cells (Fig. 2*A*, *B* and Fig S2). In contrast, PIPKIγ, PIPKIIα, and PIPKIIβ, which also contribute to the nuclear PIP_2_ pool (21), interacted minimally or not with MDM2 (Fig. 2*C* and Fig. S2). However, IPMK which converts PIP_2_ to PIP_3_, PTEN, which dephosphorylates PIP_3,_ and phosphatidylinositol transfer protein β isoform (PITPβ), which transfers PI to p53 to enable PI4P linkage (4, 22), bound MDM2 to varying degrees (Fig 2*C*, and Fig, S2), consistent with the observed interaction of PI4P and PIP_3_ with MDM2 (Fig. 1*A*).

**Fig. 2:**
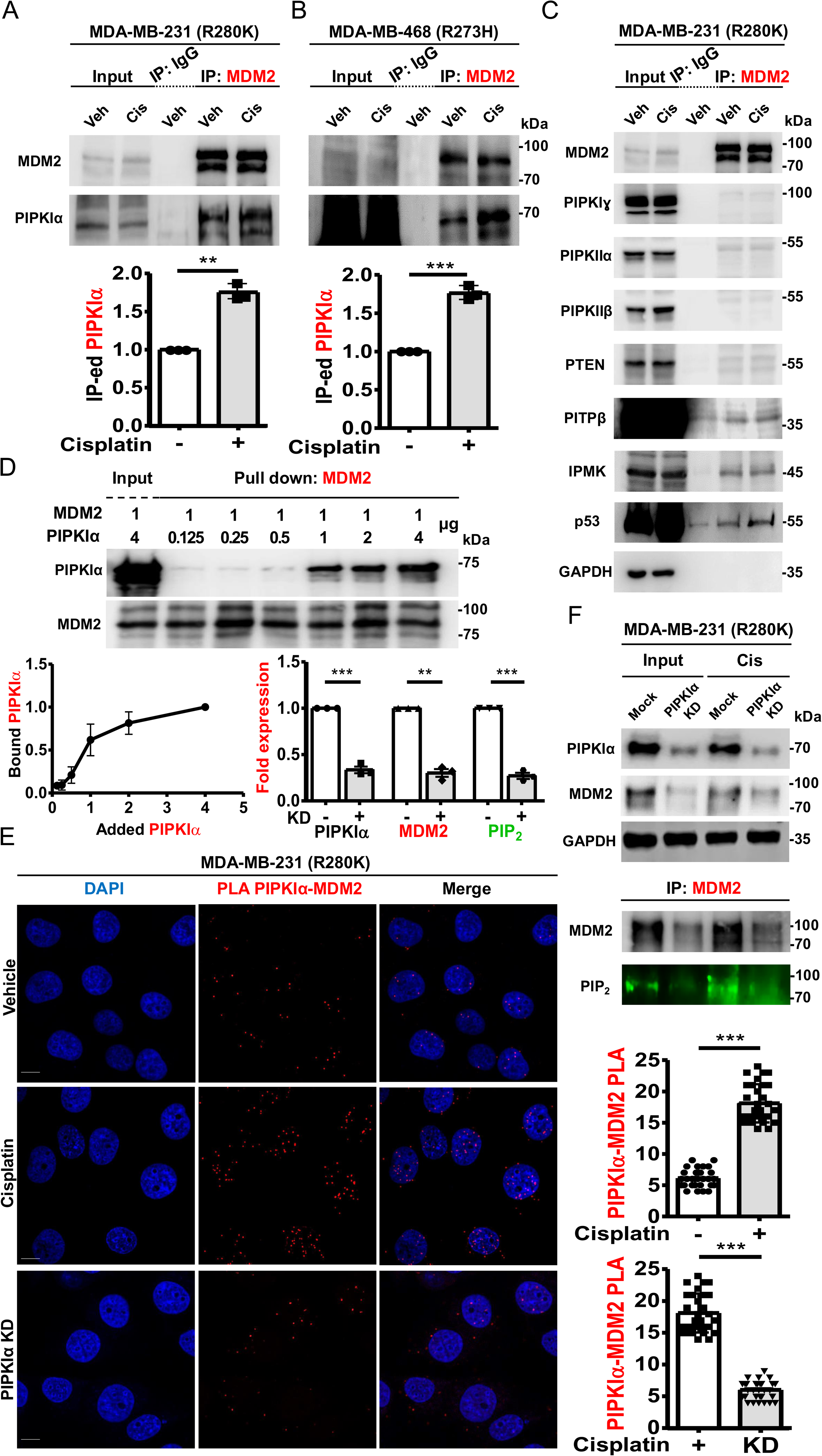
PIPKIα interacts with MDM2 and regulates PIP_2_ association. ***A, B*,** Co-IP of MDM2 with PIPKIα from MDA-MB-231 (*A*) and MDA-MB-468 cells (*B*) treated with 30LµM cisplatin or vehicle for 24Lh. Data shown represent three independent experiments. The PIPKIα IB intensity was quantified, and the graph is shown as meanL±Ls.d. of nL=L3 independent experiments. Veh, vehicle; Cis, cisplatin-treated. ***C*,** Co-IP of endogenous MDM2 from MDA-MB-231 cells treated with vehicle or 30LµM cisplatin for 24Lh. The MDM2, PIPKIα, PIPKIγ, PIPKIIα, PIPKIIβ, PTEN, PITPβ, IPMK, p53 and GAPDH IPed by MDM2 were analyzed by IB. ***D*,** Recombinant MDM2 protein (1 μg) was incubated with 0.125, 0.25, 0.5, 1, 2 or 4 μg of PIPKIα protein. MDM2 was pulled down, and the binding with PIPKIα was analyzed with an anti-PIPKIα antibody. The graphs are shown as meanL±Ls.d. of nL=L3 independent experiments. ***E*,** Quantification of nuclear PLA of PIPKIα-MDM2 in MDA-MB-231 cells treated with vehicle or 30LµM cisplatin for 24Lh. nL=L30 cells pooled from 3 independent experiments, 10 cells per experiment. ***F*,** MDA-MB-231 cells were transfected with siRNAs targeting PIPKIα. After 24 h of transfection, cells were treated with 30LµM cisplatin for 24Lh. IB analyzed the expression of the indicated proteins, IBs were quantified, and the graph is shown as meanL±Ls.d. of nL=L3 independent experiments. Mocm, empty vector; KD, knockdown; Cis, cisplatin-treated.

To elucidate whether PIPKIα binds directly to MDM2, a fixed concentration of recombinant MDM2 was incubated with increasing amounts of recombinant PIPKIα and pulled down with anti-MDM2 agarose. Immunoblotting revealed saturable binding between PIPKIα and MDM2 (Fig. 2*D*). MST confirmed a direct high-affinity interaction between PIPKIα and MDM2 (K_d_ 28±3 nM, Table 1*A*). PLA demonstrated that PIPKIα associated with MDM2 predominantly in the nucleus and that these complexes were increased by genotoxic stress and decreased by PIPKIα knockdown (KD, Fig. 2*E*). Notably, PIPKIα KD also reduced MDM2 levels and PIP_2_ association with MDM2 under basal conditions and in response to stress (Fig. 2*F* and Fig. S3*A, B*). These results indicate that PIPKIα binds MDM2 in the nucleus and is required to generate nuclear MDM2-PIP_2_ complexes.

### MDM2 associates with sHSPs in the nucleus

The sHSPs, αBC and HSP27, bind p53 in the nucleus and promote the stability of p53 (3). To investigate whether sHSPs bind MDM2, a fixed concentration of recombinant MDM2 protein was incubated with increasing amounts of sHSPs, and the complex was then pulled down with anti-MDM2 agarose. Immunoblotting (IB) showed saturable interactions between the sHSPs and MDM2 (Fig. 3*A*, *B*). High-affinity interactions between MDM2 and αBC (K_d_ 58±5 nM) and HSP27 (K_d_ 50±7 nM) were confirmed by MST (Table 1*A*). Each sHSP Co-IPed with MDM2 in mutant p53, wild-type p53, or p53-null cancer cells (Fig. 3*C* and Fig. S2). IF staining revealed that αBC and HSP27 translocate to the nucleus in response to genotoxic stress consistent with prior reports (23–25) and co-localize with MDM2 in the nucleus (Fig. 3*D*, *E*). Stress increased the formation of MDM2-αBC and MDM2-HSP27 complexes in the nucleus as determined by PLA (Fig 3*F*, *G*). These data demonstrate that αBC and HSP27 directly bind MDM2 in the nucleus in response to genotoxic stress.

**Fig. 3:**
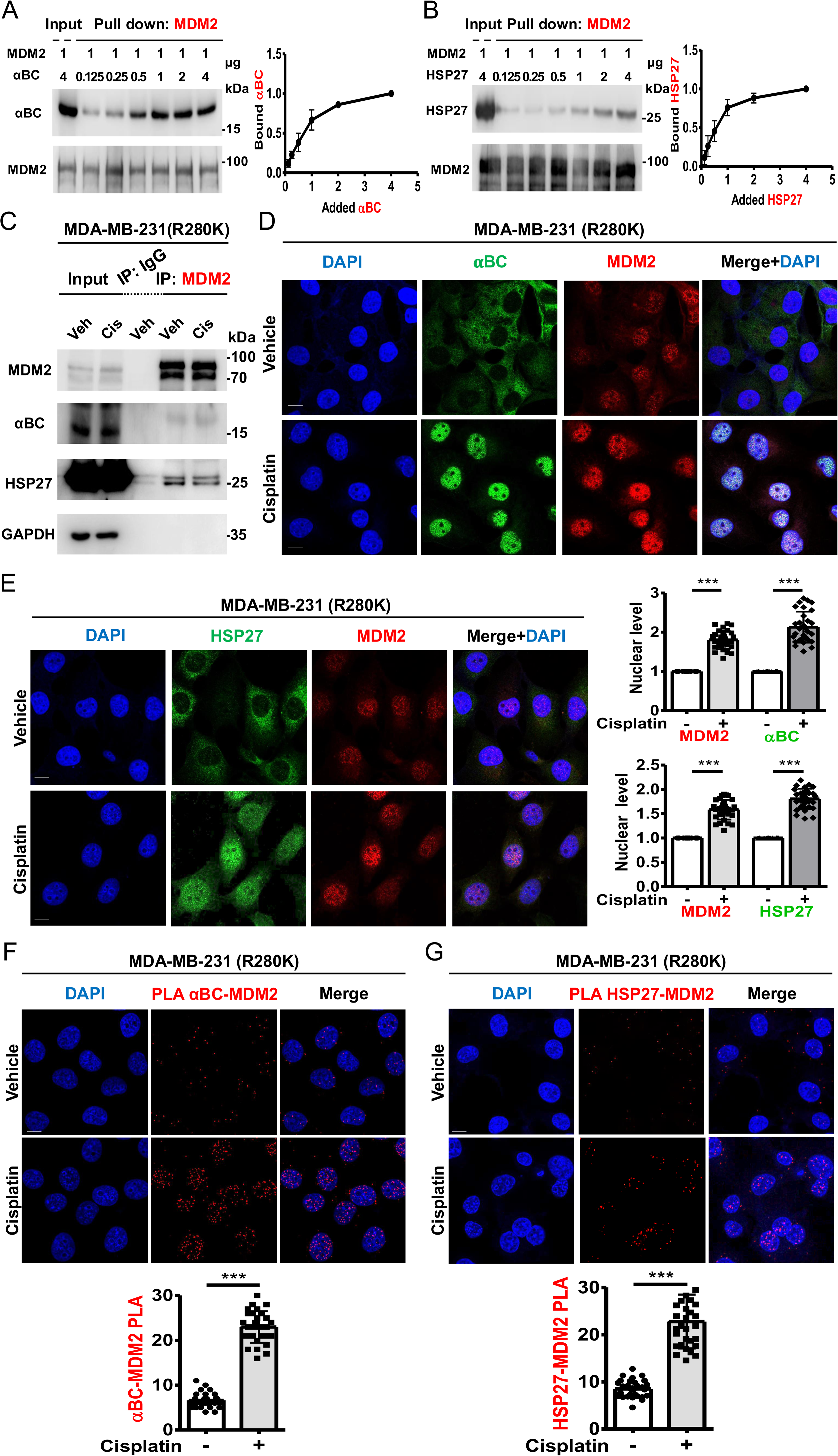
MDM2 associates with sHSPs in the nucleus. ***A*, *B*,** Recombinant MDM2 protein (1 μg) was incubated with 0.125, 0.25, 0.5, 1, 2, or 4 μg of αBC (*A*) and HSP27 (*B*) proteins. MDM2 was pulled down, and the association with αBC and HSP27 was analyzed using anti-αBC and anti-HSP27 antibodies. The graphs are shown as meanL±Ls.d. of nL=L3 independent experiments. ***C*,** Co-IP of endogenous MDM2 from MDA-MB-231 cells treated with vehicle or 30LµM cisplatin for 24Lh. IB analyzed the MDM2, αBC, HSP27, and GAPDH IPed by MDM2. Veh, vehicle; Cis, cisplatin-treated. ***D*, *E*,** Confocal images of IF staining against αBC (*D*) and HSP27 (*E*) along with MDM2 in MDA-MB-231 cells treated with vehicle or 30LµM cisplatin for 24Lh. The nuclear levels of MDM2, αBC, and HSP27 were normalized to vehicle-treated cells, and their nuclear co-localization was determined by immunofluorescence. nL=L30 cells pooled from 3 independent experiments, 10 cells per experiment. ***F*, *G*,** PLA of αBC–MDM2 (*F*) and HSP27-MDM2 (*G*) in MDA-MB-231 cells treated with vehicle or 30LµM cisplatin for 24Lh. The nuclear PLA foci of αBC–MDM2 and HSP27-MDM2 were quantified. nL=L30 cells pooled from 3 independent experiments, 10 cells per experiment.

### PIP_2_ regulates the interaction of MDM2, sHSPs and p53

To examine whether PIP_2_ regulates the association of sHSPs with MDM2 as previously demonstrated for sHSPs and p53 (3), increasing amounts of PIP_2_ were added to recombinant sHSPs and MDM2, and the complex was then pulled down with anti-MDM2 agarose. Intriguingly, while PIP_2_ enhanced the binding of both αBC and HSP27 to p53 (3), it differentially regulated sHSPs binding to MDM2 (Fig. 4*A*, *B*). Specifically, PIP_2_ increased αBC binding to MDM2 but decreased HSP27 binding to MDM2. These data suggest that αBC and HSP27 may have different roles in regulating MDM2. To investigate this possibility, αBC and HSP27 were each KDed, and MDM2 levels were assessed (Fig 4*C*). αBC KD modestly decreased MDM2 levels, while HSP27 KD had little effect on MDM2 levels (Fig. 4*C* and Fig. S3*C*). Since PIP_2_ enhances αBC binding to MDM2 and increases MDM2 protein levels, we postulated that αBC promotes MDM2 stability by inhibiting its degradation by the ubiquitin-proteasome pathway. Consistent with this idea, the proteasome inhibitor MG132 attenuated the effects of αBC KD on MDM2 levels (Fig. 4*D*). These data indicate that PIP_2_ differentially regulates sHSPs binding to MDM2 and that αBC regulates MDM2 stability by blocking proteasome degradation.

**Fig. 4:**
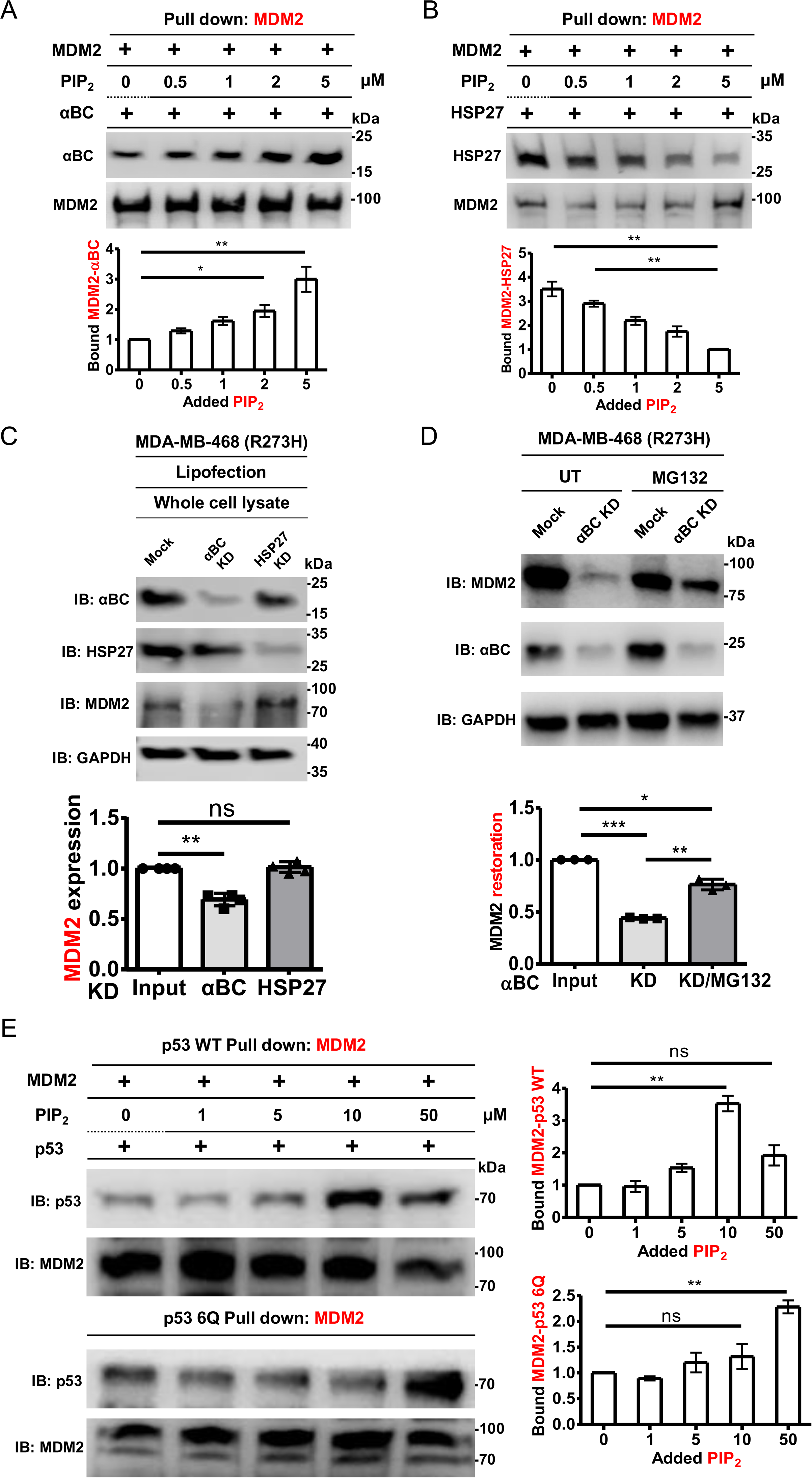
PIP_2_ regulates the interaction of MDM2, sHSPs and p53 and controls MDM2 stability. ***A*,*B*,** 0.1Lμg of MDM2 recombinant protein and 0.1Lμg of αBC (*A*) or HSP27 (*B*) protein were incubated with 0, 0.5, 1, 2, or 5LμM PIP_2_. MDM2 was pulled down, and the associated αBC or HSP27 was analyzed with an anti-αBC or an anti-HSP27 antibody. The graphs are shown as meanL±Ls.d. of *n*L=L3 independent experiments. ***C***, MDA-MB-468 cells were transfected with siRNAs for αBC or HSP27 for 48 h. Empty vector (Mock) was used as a negative control. Expression of the indicated proteins was analyzed by IB, MDM2 IBs were quantified, and the graph is shown as meanL±Ls.d. of nL=L4 independent experiments. KD, knockdown. ***D***, MDA-MB-468 cells were transfected with siRNAs for αBC for 48 h. Before harvesting, cells were treated with 10 μM of MG132 for 4 h. Empty vector (Mock) was used as a negative control. Expression of the indicated proteins was analyzed by IB, and MDM2 IBs were quantified. The graph is shown as meanL±Ls.d. of nL=L3 independent experiments ***E*,** Recombinant MDM2 protein (0.1Lμg) and p53 WT or PIP_2_ binding-defective p53 6Q mutant (p53 6Q, 0.1 μg) were incubated with 0, 1, 5, 10, or 50LμM PIP_2_. MDM2 was pulled down, and the associated p53 was analyzed with an anti-p53 antibody. The graphs are shown as meanL±Ls.d. of *n*L=L3 independent experiments. p53 WT, wild type p53; p53 6Q, p53 6Q mutant.

Next, we explored the effects of PIP_2_ on the MDM2-p53 interaction using an *in vitro* pull-down assay. Recombinant GST-p53 protein and MDM2 were incubated with increasing concentrations of PIP_2,_ and the amount of p53 that co-IPed with MDM2 was determined (Fig. 4*E*). PIP_2_ enhanced the binding of MDM2 to p53 with maximal effects observed at 10 mM PIP_2_. Since both MDM2 and p53 associate with PIP_2_, we used a PIP_2_ binding-defective p53 6Q mutant (p53 6Q), which has five lysine residues and one arginine in the p53 CTD mutated to glutamine. Notably, the incubation of PIP_2_ with MDM2 and p53 6Q resulted in the enhanced binding of MDM2 to p53, indicating that PIP_2_ binding to MDM2 explicitly increases its interaction with p53. Ectopically expressed PIP_2_ binding-defective p53 mutants (6Q and p53 R379Q mutant) in p53-null H1299 cells also bound to MDM2 consistent with prior results (Fig. S4*C*) (3). The findings indicate that PIP_2_ regulates the interaction between MDM2 and p53, at least in part, by binding to MDM2.

To determine whether sHSPs contribute to regulating MDM2–p53 interaction by PIP_2_, we performed pull-down experiments using recombinant proteins in the presence and absence of PIP_2_. Without PIP_2_, both sHSPs had little effect on MDM2 binding to p53 (Fig. 5*A*, *B*). However, in the presence of PIP_2_, αBC enhanced the interaction between MDM2 and p53, whereas HSP27 inhibited their association. These findings suggest that sHSPs differentially regulate MDM2–p53 binding through a PIP_2_-dependent mechanism.

**Fig. 5:**
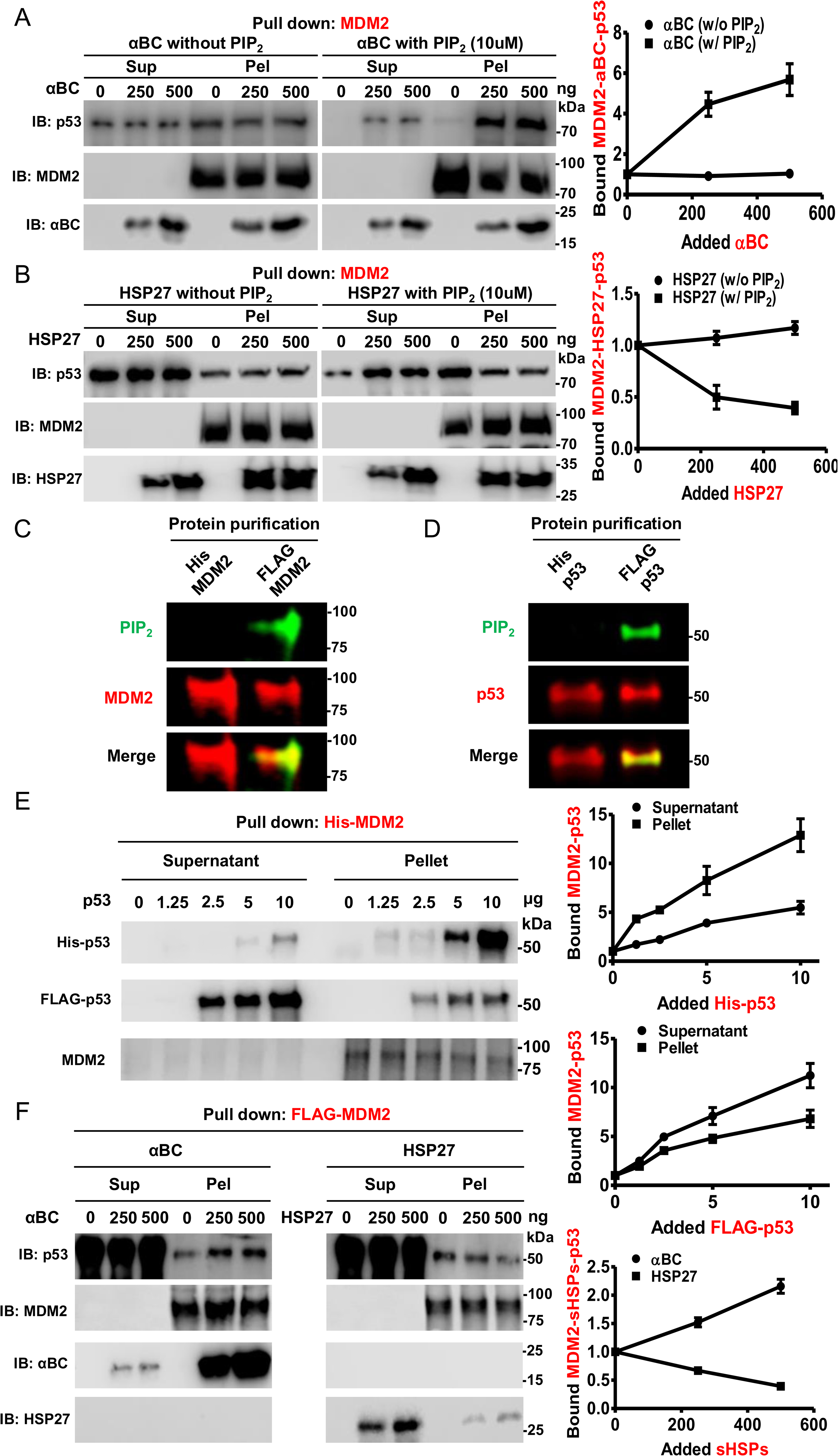
PIP_2_ and sHSPs regulate the MDM2-p53 interaction. ***A*,*B*,** 0.1 μg of MDM2 and p53 recombinant protein were incubated with gradually increasing amounts (0, 250, 500 ng) of αBC (*A*) or HSP27 (*B*) in the presence or absence of 10 μM of PIP_2_. After pulling down with MDM2 antibody-conjugated beads, the bound p53, αBC and HSP27 were analyzed with an anti-p53, anti-αBC or anti-HSP27 antibody in the supernatant and pellet. The graphs are shown as meanL±Ls.d. of *n*L=L3 independent experiments. sup, supernatant; pel, pellet. ***C***,***D***, 0.5 μg Bacteria (His-tagged) and mammalian-expressed (FLAG-tagged) MDM2 (*C*) and p53 (*D*) proteins were analyzed by fluorescence IB. Fluorescence IB detects purified proteins and PIP_2_ association. The MDM2 and PIP_2_ association is shown in the merge image. ***E***, Recombinant His-MDM2 protein (1Lμg) was incubated with 0, 1.25, 2.5, 5, or 10Lμg of His-p53 and FLAG-p53. MDM2 was pulled down, and the bound p53 was analyzed with an anti-p53 antibody in the supernatant and pellet. The graphs are shown as meanL±Ls.d. of *n*L=L3 independent experiments. ***F*,** 0.1 μg of FLAG-MDM2 and p53 recombinant protein were incubated with gradually increasing amounts (0, 250, 500 ng) of αBC or HSP27. After pulling down with MDM2 beads, the bound p53, αBC and HSP27 were analyzed with an anti-p53, anti-αBC or anti-HSP27 antibody in the supernatant and pellet. The graphs are shown as meanL±Ls.d. of *n*L=L3 independent experiments. FLAG-tagged proteins; mammalian-derived proteins.

One notable limitation of these experiments is that incubating PIP_2_ with bacteria-derived recombinant p53 and MDM2 in cell-free systems leads to binding, but PIP_n_ has rarely been found in bacteria; it is not the suitable method to demonstrate the stable association of PIP_2_ to these proteins observed in eukaryotic cellular systems (26). We purified MDM2 and p53 from mammalian HEK293FT cells to address this issue. PIP_2_ is stably associated with FLAG-tagged MDM2 and p53 produced in mammalian cells, but not His-tagged MDM2 and p53 produced in *E. coli* (Fig. 5*C*, *D*). In contrast to our *in vitro* binding assay in which PIP_2_ addition increased the association of bacterially expressed p53, MDM2, FLAG-MDM2, and FLAG-p53 exhibited reduced binding to p53 (Table 1*B*). The subsequent *in vitro* binding assay between His-MDM2, His-p53, and FLAG-p53 further supported the MST data, showing that while most of the His-p53 pulled down with MDM2 in the pellet, FLAG-p53 predominantly remained unbound in the supernatant (Fig. 5*E*). These results indicate that PIP_2_ stable association with mammalian cell-derived MDM2 and p53 inhibits the MDM2-p53 interaction.

Additionally, we analyzed the binding affinities between sHSPs and FLAG-tagged proteins. Consistent with the *in vitro* binding data (Fig. 4*A*), FLAG-MDM2 exhibited an increased binding affinity for αBC (K_d_ = 7 ± 1 nM) and a decreased affinity for HSP27 (K_d_ = 125 ± 9 nM) compared to His-MDM2 as determined by MST (Table 1*C*). In contrast, FLAG-p53 demonstrated an increased binding affinity for both αBC (K_d_ = 14 ± 1 nM) and HSP27 (K_d_ = 38 ± 3 nM). An *in vitro* binding assay involving FLAG-MDM2, FLAG-p53, and sHSPs revealed that p53 predominantly remained in the supernatant due to its association with PIP_2_. However, a small amount pulled down with MDM2 (Fig. 5*F*). Interestingly, two distinct patterns emerged: αBC enhanced the interaction between FLAG-MDM2 and p53, whereas HSP27 reduced their binding, which mimicked the results observed by *in vitro* binding assay with the addition of PIP_2_. Taken together, these data from both His- and FLAG-proteins indicate that sHSPs and stably associated versus bound PIP_2_ differentially regulate MDM2–p53 binding.

### PIP_2_ and sHSPs regulate the function of MDM2

We next examined whether sHSPs regulated the ubiquitination of MDM2. A unique characteristic of RING finger E3 ubiquitin ligases, including MDM2, is that they regulate the ubiquitination of their targets and themselves (autoubiquitination) (27). As such, we utilized an *in vitro* ubiquitination assay to determine the effects of PIP_2_ and sHSPs on MDM2 autoubiquitination. Compared to the positive control, MDM2 *in vitro* ubiquitination was decreased by αBC but increased when HSP27 and PIP_2_ were added individually (Fig. 6*A*). Notably, sHSPs regulated MDM2 *in vitro* ubiquitination in a dose-dependent manner while the effects of PIP_2_ were dose-independent. Moreover, adding PIP_2_ to this cell-free MDM2 autoubiquitination assay did not alter the impact of αBC or HSP27 on MDM2 autoubiquitination (Fig. S4*A*). These data demonstrate that PIP_2_ and sHSPs regulate MDM2 autoubiquitination, with αBC inhibiting ubiquitination and HSP27 enhancing MDM2 autoubiquitination.

**Fig. 6:**
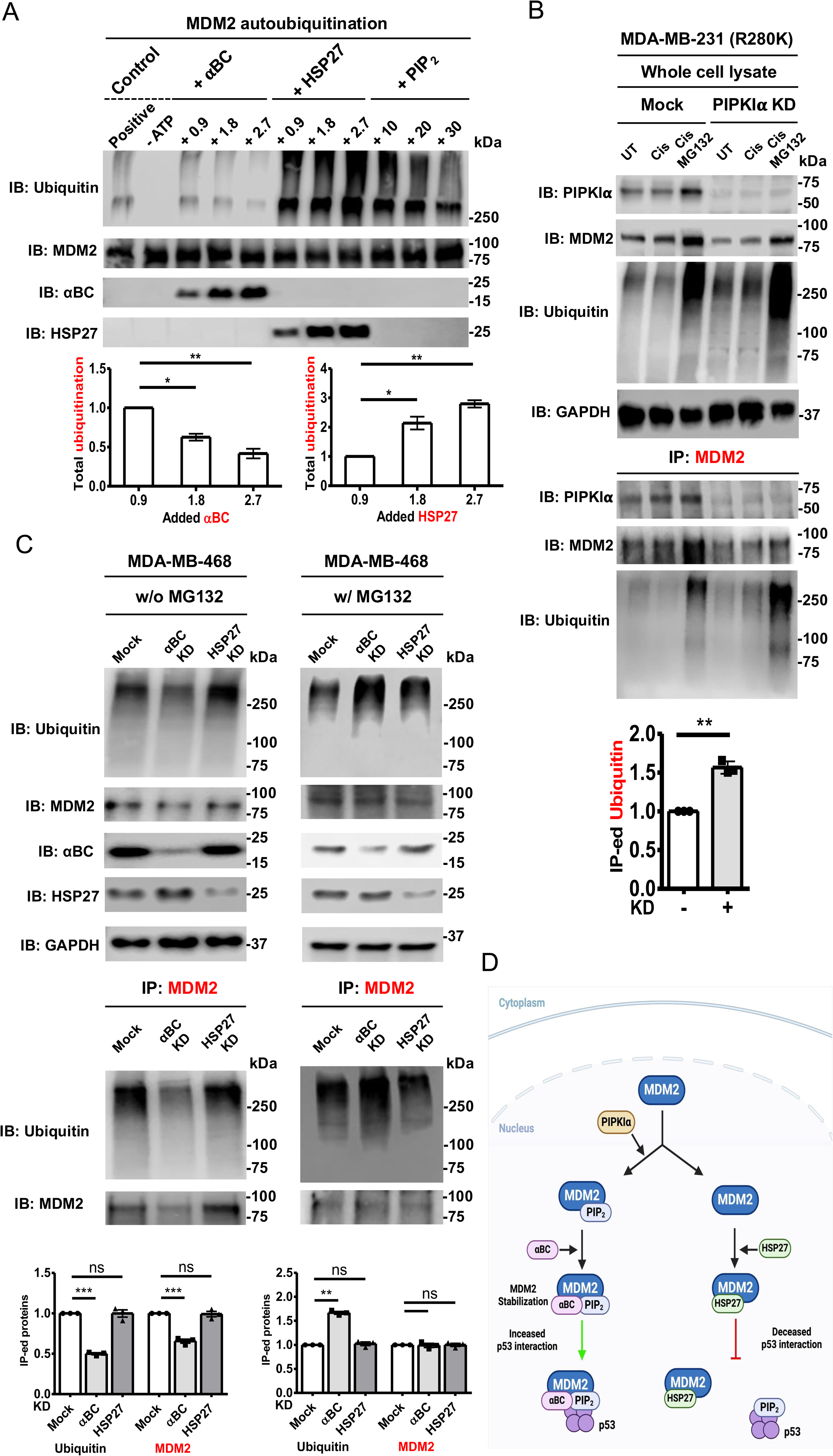
PIP_2_ and sHSPs regulate the ubiquitination function of MDM2. ***A*,** For MDM2 autoubiquitination, 100 nM of E1 enzyme, 1 μM of E2 enzyme, 1LμM of MDM2, E3 ligase reaction buffer, 10mM of MgATP solution and 100 μM of ubiquitin were incubated with different concentrations of αBC, HSP27 (0.9, 1.8, or 2.7 μM), or PIP_2_ (10, 20, or 30 μM) for 1 h. Ubiquitin, MDM2, αBC and HSP27 were analyzed by IB, and ubiquitin IBs were quantified. The graphs are shown as meanL±Ls.d. of *n*L=L3 independent experiments. ***B***, MDA-MB-231 cells were transfected with siRNAs for PIPKIα and treated with vehicle or 30LµM cisplatin for 24Lh. Empty vector (Mock) was used as a negative control. After cells were treated with 10 μM of MG132 for 4 h, cells were harvested for IP of MDM2. Expression of the indicated proteins was analyzed by IB, and ubiquitin IBs were quantified. The graph is shown as meanL±Ls.d. of nL=L3 independent experiments. KD, knockdown; UT, untreated; Cis, cisplatin treated; Cis/MG132, Cisplatin/MG132 treated. ***C*,** MDA-MB-468 cells were transfected with siRNAs for αBC and HSP27 for 48 h in absence or presence of MG132. Empty vector (Mock) was used as a negative control. Cells were processed for IP of MDM2. IB was used to analyze indicated proteins, and ubiquitin IBs were quantified. The graphs are shown as meanL±Ls.d. of nL=L3 independent experiments. KD, knockdown. **D**, A model of PIP_2_ regulation of the interaction between MDM2 and sHSPs, which differentially control the stability of MDM2 and interaction of MDM2 and p53. In the presence of PIPKIα and PIP_2_, MDM2 binds αBC, resulting in the stabilization of MDM2 and enhanced MDM2-p53 interaction. Conversely, in the absence of PIPKIα and PIP_2_, HSP27 is recruited to MDM2, which inhibits the interaction of MDM2 with p53.

Next, we investigated the regulation of MDM2 ubiquitination in cellular models. MDM2 was IPed after PIPKIα KD in the presence or absence of Cisplatin and MG132. Immunoblotting showed the successful KD of PIPKIα, increasing MDM2 ubiquitination (Fig. 6*B*). Along with the increased accumulation of ubiquitinated MDM2, MDM2 expression was decreased and restored by MG132 treatment. Similar results were observed in A549 cells expressing wild-type p53 (Fig S4*B*). These data indicate that KD of PIPKIα and the resulting decrease in PIP_2_ enhances the ubiquitination of MDM2, leading to its degradation by the ubiquitin-proteasome pathway.

To identify the functional role of sHSPs in the ubiquitination of MDM2, we transfected siRNAs targeting sHSPs in p53 mutant MDA-MB-468 cells to measure degradation of ubiquitinated MDM2 in the absence of MG132 and accumulation of ubiquitinated MDM2 in the presence of MG132 (Fig. 6*C*). In the absence of MG132, αBC KD resulted in decreased accumulation of ubiquitinated MDM2 and decreased level of MDM2 due to degradation. However, following MG132 treatment, αBC KD led to an increased accumulation of ubiquitinated MDM2, and MDM2 levels were similar to that in the control group. Despite MG132 treatment, HSP27 KD did not significantly alter MDM2 ubiquitination and/or MDM2 levels. These data indicate that sHSPs have differential effects on MDM2 ubiquitination *in vitro* and in cellular models.

Previous studies have shown that MDM2 regulates several additional targets by ubiquitination, including FOXO3a and histone H2A/H2B, in a p53-independent manner (28, 29). To test whether PIP_2_ association via PIPKIα modulated targets MDM2 targets in a p53- independent system, we employed PIPKIα KD using siRNA in p53-null H1299 cell line (Fig. S4*D*). Consistent with our earlier findings, PIPKIα KD led to a reduction of MDM2 levels, resulting in stabilizing its targets, FOXO3a and H2B. Notably, no significant change was observed in H2A levels, which aligns with a previous report that H2A undergoes MDM2- mediated ubiquitination *in vitro* but not *in vivo* (28). These data imply that PIP_2_’s association with MDM2 also regulates the ubiquitination of other MDM2 substrates, expanding the functional effects of the MDM2-PIP_2_ complex.

## Discussion

Since the discovery of PIP_n_s in the nucleus (30), their function in the nucleus has been intensively investigated (5, 12, 19, 31–36). Recently, the lipid kinase PIPKIα and its product PIP_2_ were reported to bind the CTD of stress-activated wild-type p53 and mutant p53 and stabilize p53 by recruiting the sHSPs αBC and HSP27 to the p53-PIP_2_ complex^3^. Since the stability of wild-type p53 is regulated by E3 ubiquitin ligase MDM2 (37), we began investigating its potential functional role in regulating the effects of PIP_2_ on p53 stability. We discovered that multiple PIP_n_s interact with MDM2. Specifically, to a lesser extent, PIP_2,_ PI4P, and PIP_3_ bind to MDM2 in response to stress. The association of PIP_2_ with MDM2 requires PIPKIα, which binds directly to MDM2 with high affinity (K_d_ 28±3 nM) to transfer its product PIP_2_ to MDM2, a mechanism highly reminiscent of their role in regulating p53 stability (3). The association between MDM2 and PIP_2_ is very stable as it can be detected by IB and withstands harsh denaturation with reducing agents, high temperature, and SDS-polyacrylamide gel electrophoresis, similar to other PIP_2_ effectors (3, 38, 39). Moreover, metabolic labeling of cells with [H^3^]*myo*-inositol, which is metabolized solely to phosphoinositides, results in the stable incorporation of [H^3^] label into MDM2 in response to stress, and this linked [H^3^]PIP_n_ is also resistant to harsh denaturing conditions. These results indicate that PIP_2_ and other PIP_n_s are closely related to MDM2, likely by a covalent bond/novel posttranslational modification (PTM). The chemical nature and location of this putative PTM on MDM2 and p53 are currently being investigated by multiple orthogonal approaches.

Once PIP_2_ interacts with MDM2, it regulates the association of MDM2 with sHSPs. As previously reported, genotoxic stress induces the nuclear translocation of sHSPs (3, 23, 25, 40, 41), facilitating their interaction with nuclear MDM2. In contrast to p53, where PIP_2_ binding enhances the recruitment of αBC and HSP27, which both stabilize p53 (3), the PIP_2_ bound to MDM2 differentially regulates αBC and HSP27 recruitment, acting like a switch (Fig. 6*D*). Specifically, PIP_2_ binding to MDM2 enhances αBC binding, which stabilizes MDM2. In contrast, the association of PIP_2_ with MDM2 decreases HSP27 binding, which does not affect MDM2 stability. Moreover, recombinant αBC inhibits MDM2 autoubiquitination *in vitro*, while HSP27 enhances MDM2 autoubiquitination. To add to the complexity of these interactions, αBC and HSP27 also have divergent effects on the MDM2-p53 interaction, with αBC increasing the association of MDM2 with p53 by a PIP_2_-dependent mechanism and HSP27 having the opposite effect. These opposing effects of αBC and HSP27 enable exquisite fine-tuning of the regulation of the MDM2-p53 nexus by PIP_2_.

Since adding PIP_2_ to the *in vitro* binding assay incubation does not fully recapitulate the stable association of PIP_2_ to p53 and MDM2 observed in cells, we purified PIP_2_-associated MDM2 and p53 protein from mammalian HEK293FT cells, notably, unlike bacteria-derived MDM2 and p53, mammalian-derived MDM2 and p53 with stably associated PIP_2_ exhibit reduced interaction. Despite these differences, mammalian-expressed MDM2 showed increased αBC interaction and diminished HSP27 association, consistent with our findings in cell-free studies. Moreover, αBC enhanced the interaction between MDM2 and p53, while HSP27 inhibited their interaction, consistent with the cell-free results. These data point to a complex, highly tunable system whereby PIP_2_ differentially regulates sHSPs’ interactions with MDM2 with distinct functional consequences regarding MDM2 autoubiquitination and p53 interaction and stability.

Since the interaction between MDM2 and p53 is critical for p53 stability and oncogenic activity (42–44), our results unveil a new therapeutic target: the MDM2-PIP_2_-sHSPs complex, which controls the MDM2-p53 interaction. Importantly, PIP_2_ selectively regulates sHSP interactions with MDM2 and p53 to control the stability of MDM2. While studies have investigated how sHSPs promote the ubiquitination of diverse targets (45–49), no reports have demonstrated that sHSPs regulate MDM2 ubiquitination. Our data point to a model that nuclear PIP_2_ acts like an on-off switch that confers specificity to determine whether αBC or HSP27 are recruited to MDM2, resulting in opposing effects (Fig. 6*D*). Through this association, it regulates MDM2 stability, downstream targets, ubiquitination function, and interaction with p53. In addition, our results provide additional support for a novel PIP_n_ signaling pathway on proteins such as p53 (3, 12), the nuclear poly(A) polymerase Star-PAP (5), and MDM2, where they function as ‘third messengers’ tightly linked to effector proteins to regulate their stability and function.

## Experimental procedures

### Cell culture and constructs

MDA-MB-231, MDA-MB-468, BT-20, A549, HCT116, H1299, and HEK293FT cells were purchased from ATCC (American Type Culture Collection). BT-20 cells were maintained in EMEM (#30-2003, ATCC) supplemented with 10% fetal bovine serum (#SH30910.03, Hyclone) and 1% penicillin/streptomycin (#15140-122, Gibco). All other cells were maintained in DMEM (#11995-065, Gibco) with the abovementioned supplements. All cell lines were routinely checked for mycoplasma contamination using the MycoAlert Mycoplasma Detection Kit (#LT07-318, Lonza), and only mycoplasma-negative cells were used. None of the cell lines are listed in the database of commonly misidentified cell lines maintained by the International Cell Line Authentication Committee (ICLAC). The FLAG-tagged wild-type p53, p53 6Q mutant, p53 R175H mutant, αBC and HSP27 constructs were described previously(3). The MDM2 cDNA was purchased from OriGene. Constructs were transfected into mammalian cells using a lipid-based delivery system, Lipofectamine^TM^3000 (#L3000015, Thermo Fisher Scientific), according to the manufacturer’s instructions. Typically, 2–5 μg DNA and 6–10 μl lipid were used for transfections in 6-well plates. Transfection efficiency was determined by immunoblotting. For recombinant protein production, the His-tagged constructs for MDM2, αBC, HSP27, GST-tagged p53 6Q mutant and PIPKIα were purchased from Genscript and the recombinant proteins were purified as previously described(3). Recombinant GST-tagged p53 (#14-865) was purchased from MilliporeSigma.

### Antibodies and reagents

Monoclonal antibodies against PIP_2_ (clone 2C11, #Z-P045, Echelon Biosciences), PIP_2_ (clone KT10, #MSBS2283, MilliporeSigma), p53 (clone DO-1, #SC-126, Santa Cruz Biotechnology), HSP27 (clone F-4, #SC-13132, Santa Cruz Biotechnology), αBC (D6S9E, #45844, Cell Signaling), PIPKIIα (PIP4K2A, clone D83C1, #5527, Cell Signaling), PTEN (clone D4,3, #9188, Cell Signaling), GAPDH (clone 0411, #SC-47724, Santa Cruz Biotechnology), and polyclonal antibodies against MDM2 (clone D1V2Z, #86934, Cell Signaling), MDM2 (#AF1244, R&D Systems), αBC (#ab13497, Abcam), PIPKIα (PIP5K1A, #9693, Cell Signaling), PIPKIγ (PIP5K1C, #3296, Cell Signaling), PIPKIIβ (PIP4K2B, #9694, Cell Signaling), PITPNB (#NBP2-19841, Novus Biologicals), FOXO3a (clone 75D8, #2497, Cell Signaling), Histone H2A (#2578, Cell Signaling), Histone H2B (clone D2H6, #12364, Cell Signaling), Ubiquitin (P37, #58395, Cell Signaling) were utilized in this study. Polyclonal antibodies against IPMK were produced as described previously (50). For conventional immunostaining and PLA analysis of phosphoinositides, anti-PIP_2_ (#Z-P045) antibodies were used. For immunostaining analyses and proximity ligation assay (PLA), antibodies were diluted at a 1:100 ratio. The ON-TARGETplus siRNA SMARTpool with 4 siRNAs in combination against human MDM2, αBC, HSP27 and PIPKIα were purchased from Dharmacon. As a control, non-targeting siRNA was purchased from Dharmacon. RNAiMAX reagent (#13778150, Thermo Fisher Scientific) was utilized to deliver siRNAs into the cells, and knockdown efficiency was determined by immunoblotting. Cisplatin (#S1166) was purchased from Sellekchem.

### Immunoprecipitation and immunoblotting

Cells were washed once with ice-cold PBS (#14190-144, Gibco), and lysed in ice-cold RIPA lysis buffer system (#sc-24948, Santa Cruz Biotechnology) supplemented with 1 mM Na_3_VO_4_, 5 mM NaF and 1x protease inhibitor cocktail (#11836153001, Roche). The cell lysates were sonicated at 16% amplitude for 5-10 s. After sonication, the cell lysates were incubated at 4°C with continuous rotation for 30 min for lysis. Subsequently, lysates were centrifuged at maximum speed for 15 min to collect the supernatant. The protein concentration was measured by the Bradford protein assay (#5000201, BIO-RAD) and equal amounts of protein were loaded in each lane. For immunoblotting analyses, antibodies were diluted at a 1:1,000 ratio except for p53 (clone DO-1, 1:5,000) and GAPDH (clone 0411, 1:5,000). For immunoprecipitation, cell lysates were incubated with 20 µl anti- MDM2 (SMP14, #SC-965 AC, Santa Cruz Biotechnology) mouse monoclonal IgG antibody-conjugated agarose at 4°C for 24 h. Normal immunoglobulin (IgG)-conjugated agarose was used as a negative control (#sc-2343, Santa Cruz Biotechnology). After incubation, samples were washed three times with PBST (PBS with 0.05% Tween 20), the precipitated protein complex was resuspended with SDS sample loading buffer. For immunoblotting, 5-20 µg of protein were loaded. Horseradish peroxidase (HRP)- conjugated antibodies were used for immunoblotting of immunoprecipitated complexes to avoid non-specific detection of immunoglobulin in the immunoprecipitated samples. HRP-conjugated p53 (#SC-126 HRP), HSP27 (clone F-4, #SC-13132 HRP) and GAPDH (clone 0411, #SC-47724 HRP) antibodies were purchased from Santa Cruz Biotechnology. Immunoblots were captured using the Odyssey Imaging System (LI-COR Biosciences), and the intensity of protein bands was quantified using ImageJ. The unsaturated exposure of immunoblot images was used for quantification with the appropriate loading controls as standards.

### Fluorescent IP-IB

Cells were lysed and immunoprecipitated as described above. The sample was then boiled at 95°C for 10 min. For immunoblotting, 5-20 µg of protein were loaded. The protein complexes associated with MDM2 were resolved by SDS-PAGE and transferred onto a PVDF membrane (#IPVH00010, MilliporeSigma). The membrane was blocked with 3% BSA in PBS for 1 h at room temperature. For double fluorescent IP-IB detecting MDM2-PIP_2_ complex, anti-MDM2 rabbit monoclonal IgG antibody (clone D1V2Z, #86934, Cell Signaling) at 1:1,000 dilution and anti-PIP_2_ mouse monoclonal IgM antibody (#Z-P045, Echelon Biosciences) at 1:1,000 dilution were incubated together in blocking buffer and incubated with the membrane at 4°C overnight. Then the membrane was washed three times with PBST for 10 min each time on the rocking incubator. For the secondary antibody incubation, goat anti-rabbit IgG antibody conjugated with IRDye 800CW fluorophore (#926-32211, LI-COR) detectable on the 800 nm wavelength channel of the Odyssey Fc Imaging System (LI-COR Biosciences) and goat anti-mouse IgM antibody conjugated with IRDye 680RD fluorophore (#926-68180, LI-COR) detectable on the 700 nm wavelength channel at 1:5,000 dilution were mixed together in blocking buffer and incubated with the membrane at room temperature for 2 h. The membrane was then washed three times with PBST for 10 min each time on the rocking incubator. The images were subsequently acquired using the 700 and 800 nm wavelength channels simultaneously on the Odyssey Fc Imaging System (LI-COR Biosciences). The MDM2-associated PIP_2_ complex was visualized by overlapping the 700 and 800 nm channels.

### Radio-detection of ^3^H-inositol metabolic labeling

Cells were seeded and maintained in an Opti-MEM (#31985070, Thermo Fisher Scientific) supplemented with 10% dialyzed FBS with a 10,000 molecular weight cut-off (#F0392, MilliporeSigma) and 1% Pen/strep (Cat#: 15140-122, Gibco). The cells were allowed to adhere overnight to reach 5-10% confluency the next day and were treated with 25 μCi/ml of *myo*-[2-^3^H]-inositol (#NET1156005MD, PerkinElmer Life Science). Normal myo-inositol was supplemented in the medium of the control group. When the cells reached 50% confluency, they were transfected with MDM2 constructs as described above for 24 h, followed by treatment with vehicle control or 30 μM Cisplatin. After 24 h treatment, the cells were processed for IP-IB as described above. Cell lysates were subjected to IB for Coomassie blue staining and immunoblotting as described previously. For Coomassie blue staining, after running SDS-polyacrylamide gel electrophoresis, gels were washed with distilled water and stained with Coomassie blue (#ab119211, Abcam) for 1 h. Subsequently, the gels were dissected into 1∼2 mm pieces for LSC. The dissected gels were dissolved in 500 µl of 30 % hydrogen peroxide (#H1009, MilliporeSigma) at 55 °C overnight. After complete dissolution, the sliced gels were cooled down at room temperature. The dissolved samples were mixed with 10 ml of Hionic-Fluor (#6013319, PerkinElmer Life Science) in Glass Scintillation Vials (#50212992, Thermo Fisher Scientific). The samples were placed in a dark room for 30 min for dark adaptation then were loaded into Tri-Crab Scintillation counter (#4910TR PerkinElmer Life Science). Disintegration per minute (DPM) was automatically generated by Perkin Elmer Quantasmart^TM^.

### *In vitro* binding assay

Recombinant proteins were expressed in BL-21(DE3) *Escherichia coli* strain (#EC0114, Thermo Fisher Scientific). Cells were lysed by sonication in 1% Brij58. GST-tagged p53 6Q mutant was purified with GST Sepharose 4B (#17075604, Cytiva), while His-tagged MDM2, αBC, HSP27, and PIPKIα were purified with Ni-NTA-agarose (#166038887,, Qiagen) as previously described (3). Recombinant FLAG-tagged p53 R248Q were expressed in HEK293FT cells and purified protein by Meng S. Choy and Wolfgang Peti, University of Connecticut, Farmington, CT, USA. FLAG-tagged MDM2 were expressed in HEK293FT cells. Cells were lysed, and proteins were purified using FLAG ^®^ Purification Kit (CELLMM2-1KT, MilliporeSigma). After purification, eluates were dialysis to exchange buffer into PBS using a dialysis cassette (#66380, Thermo Fisher Scientific), flash-frozen, and stored at −80 °C. Recombinant GST-tagged p53 (#14-865) was purchased from MilliporeSigma. The synthetic PIP_2_ diC16 (#P-4516) was purchased from Echelon Biosciences. The binding assay was conducted in Tris-based MST buffer containing 50 mM Tris-HCl, pH 8.0, 50 mM NaCl, 80 mM KCl, and 0.05% Tween-20. Various recombinant proteins were incubated at different concentrations. For pull-down experiments, 20 μl anti-MDM2 antibody-conjugated agaroses (SMP14, #SC-965 AC, Santa Cruz Biotechnology) were used. Following overnight incubation at 4°C, unbound proteins were removed by washing three times with MST buffer or collected for supernatant and pellet comparison. The protein complex was analyzed by immunoblotting. GraphPad Prism generated the quantitative graph. The images were processed using ImageJ.

### Liposome sedimentation assay

For the sedimentation assay, control (P-B000), PI-coated (P-B001), PI4P-coated (P-B004a), PIP_2_-coated (P-B045a), and PIP_3_-coated (P-B345a) beads were purchased from Echelon Biosciences. Recombinant MDM2 was purified as described above. The sedimentation assay was performed in Tris-based MST buffer containing 50 mM Tris-HCl, pH 8.0, 50 mM NaCl, 80 mM KCl, and 0.05% Tween-20 by incubating 0.1 μg MDM2 recombinant protein with 10 μl lipid-coated beads. After overnight incubation at 4°C, unbound MDM2 was removed by washing three times with MST buffer, and the sedimentations were resuspended with an equal amount of SDS sample buffer as supernatant. Subsequently, samples were resolved by SDS–PAGE and proteins associated with the liposomes were detected by immunoblotting.

### Microscale Thermophoresis (MST) Assay

The MST assay was utilized to measure the binding affinity of purified recombinant proteins *in vitro, as described previously*(*12*). The target protein was fluorescently labeled using the Monolith Protein Labeling Kit RED-NHS 2nd Generation (#MO-L011, Nano Temper). A sequential titration of unlabeled ligand proteins, PI-PolyPIPosomes, or PI-micelles was made in a Tris-based MST buffer containing 50 mM Tris-HCl, pH 8.0, 50 mM NaCl, 80 mM KCl, and 0.05% Tween-20. This was mixed and loaded into Monolith NT.115 Series capillaries (#MO-K022, Nano Temper), and MST traces were recorded using Monolith NT.115 pico. The binding affinity was automatically generated by MO. Control v1.6 software.

### Immunofluorescence (IF) and Confocal Microscopy

For immunofluorescence studies, cells were cultured on coverslips coated with 0.2 % gelatin (#G9391, MilliporeSigma). The cells were fixed with 4% paraformaldehyde (PFA) (#sc-281692, Santa Cruz Biotechnology) for 20 min at room temperature, followed by three times washing with PBS. Then, the cells were permeabilized with 0.3% Triton-X100 for 10 min and rewashed three times with PBS. Subsequently, the cells were blocked with 3% BSA in PBS for 1 h at room temperature. After blocking, the cells were incubated with primary antibodies overnight at 4 °C.

Following primary antibody incubation, the cells were washed three times with PBS and then incubated with fluorescent-conjugated secondary antibodies (Molecular Probes) for 1 h at room temperature. Following secondary antibody incubation, the cells were washed three times with PBS, and the nuclei were counterstained with 1 µg/ml 4’,6-diamidino-2-phenylindole (DAPI) (#D3571, Invitrogen) in PBS for 30 min at room temperature. The cells were rewashed three times with PBS and mounted in Prolong^TM^ Glass Antifade Mountant media (#P36984, Thermo Fisher Scientific). Leica SP8 3xSTED Super-Resolution Microscope took imaging for high-resolution confocal microscope data. The Leica SP8 3xSTED microscope was controlled by LASX software (Leica Microsystems). All images were acquired using a 100X objective lens (N.A. 1.4 oil). For quantification, the mean fluorescent intensity of channels in each cell was measured by LASX. GraphPad Prism generated the quantitative graph. The images were processed using ImageJ.

### Proximity Ligation Assay (PLA)

As previously described, PLA detected *in situ* protein-protein/PI interactions (12). After fixation and permeabilization, cells were blocked before incubation with primary antibodies, as in routine IF staining. The cells were then processed for PLA (#DUO92101, MilliporeSigma) according to the manufacturer’s instruction and previously published. The slides were mounted with Duolink® In Situ Mounting Medium with DAPI (#DUO82040, MilliporeSigma). PLA signals were detected using a Leica SP8 confocal microscope, which visualized the signals as discrete punctate foci and provided the intracellular localization of the complex. Quantification of nuclear PLA foci was employed using ImageJ.

### *In vitro* ubiquitination assay

For the *in vitro* ubiquitination assay targeting MDM2 autoubiquitination, 100 nM of E1 enzyme (#E304, R&D Systems), 1 μM of E2 enzyme (#E2627, R&D Systems), 1 μM of MDM2, E3 ligase reaction buffer (#B71, R&D Systems), 10mM of MgATP solution (#B20, R&D Systems) and 100 μM of ubiquitin (#U100H, R&D Systems) were incubated with different concentrations of αBC, HSP27 (0.9, 1.8, or 2.7 μM), or PIP_2_ (10, 20, or 30 μM). For p53 ubiquitination, 5 μM of p53 was included in the ubiquitination mixture. A negative control was established by replacing MgATP with ddH_2_O. Recombinant MDM2, p53, αBC, and HSP27 were purified, and PIP_2_ was purchased as described above. The ubiquitination mixture was incubated for 1 h at 37 °C, and the reaction was terminated by adding 100 mM of DTT (#R0861, Thermo Fisher Scientific). Subsequently, the ubiquitin conjugation reaction products were separated using SDS–PAGE, and Ubiquitin, MDM2, αBC, and HSP27 were detected by immunoblotting. Quantitative analysis was conducted using GraphPad Prism, and images were processed using ImageJ.

### Statistics and Reproducibility

Statistical analysis of the data was performed using GraphPad Prism and Microsoft Excel, and data from at least three different experiments were used in this study. Two-tailed unpaired *t*-tests were used for pair-wise significance unless otherwise indicated. We note that no power calculations were used. Sample sizes were determined based on previously published experiments where significant differences were observed. Each experiment was independently repeated at least three times, with the number of repeats defined in each figure legend. We used at least three independent experiments or biologically independent samples for statistical analysis.

## Data availability

Data generated in this study are contained within the article.

## Supporting information

This article contains supporting information Figs. S1-S4.

## Acknowledgments

We thank W. Peti and M. S. Choy for the mammalian-derived p53 protein and comments. We are also indebted to members of the Anderson and Cryns for their helpful discussion and review of the manuscript.

## Author Contributions

JL, MC, VLC, and RAA designed the experiments. JL, MC, and TW performed the experiments. JL, VLC, and RAA wrote the manuscript.

## Funding and additional information

This work was supported in part by the National Institutes of Health grant 1R01CA286492 (R.A.A. and V.L.C.); Department of Defense Breast Cancer Research Program grants W81XWH2110129 (V.L.C.), HT9425-23-1-0553 (V.L.C.), and HT9425-23-1-0554 (R.A.A.); a grant from the Breast Cancer Research Foundation (V.L.C.); the Shenzhen Medical Research Fund grant D2301007 (M.C.); the Shenzhen Natural Science Foundation grant JCYJ20240813094605008 (M.C.); the Guangdong Province Basic and Applied Basic Research Foundation grant 2023A1515110237 (M.C.); and the National Natural Science Foundation of China grant 32400577 (M.C.).

