## Supporting information for "Regulation of the MDM2-p53 Nexus by a Nuclear Phosphoinositide and Small Heat Shock Protein Complex"

Jeong Hyo Lee, Mo Chen, Tianmu Wen, Meng S. Choy, Wolfgang Peti, Richard A. Anderson*, and Vincent L. Cryns*

*Correspondence: Richard A. Anderson and Vincent L. Cryns

**This PDF file includes:**

Figures S1 to S4 and the accompanying figure legends.

### Supporting Information

**Figure S1**

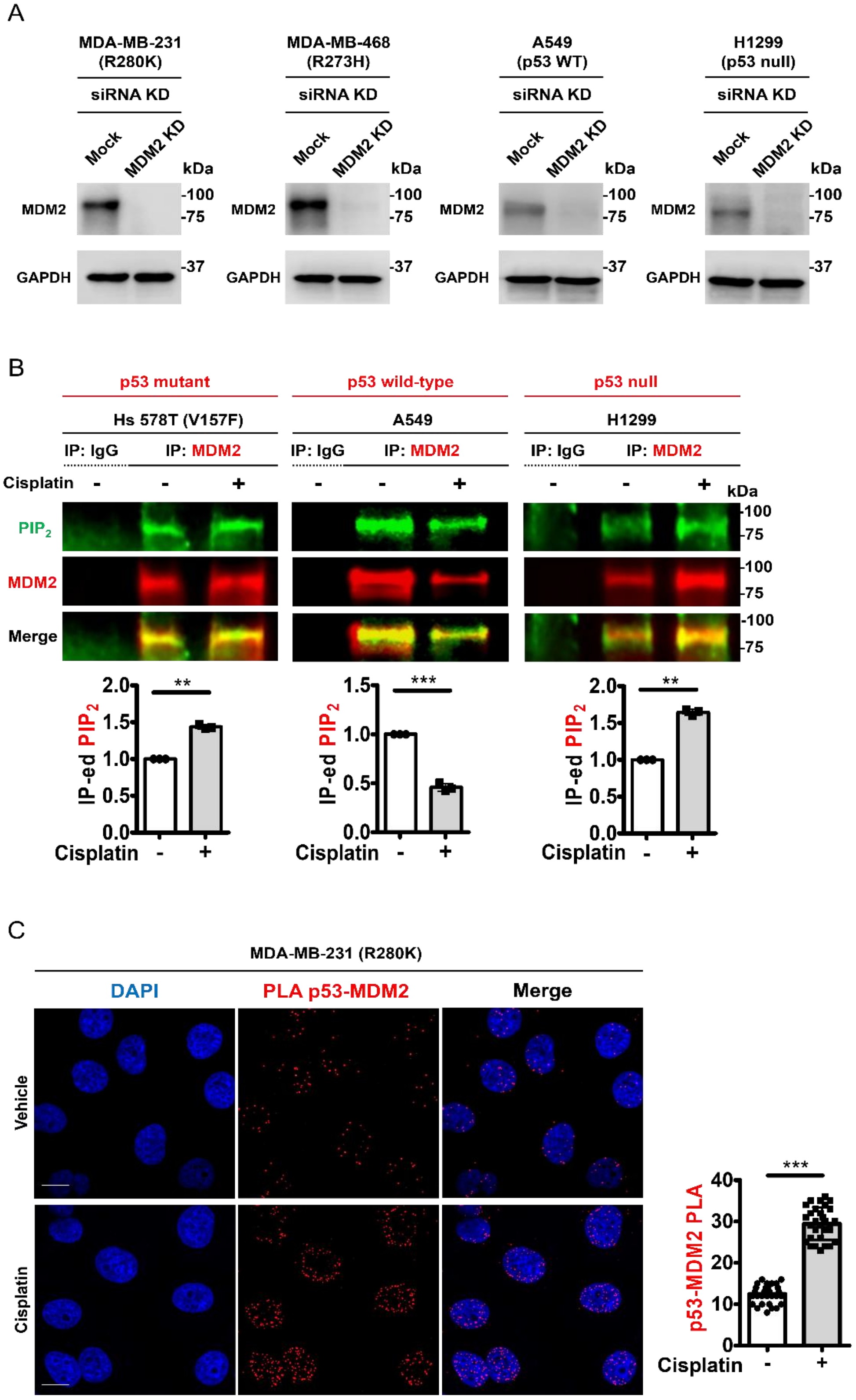

**Figure S1: MDM2 stably associates with $PIP_2$ in multiple cell lines**

***A*****,** MDA-MB-231, MDA-MB-468, A549, and H1299 cells were transfected with siRNAs targeting MDM2. After 48 h, IB analyzed endogenous MDM2. KD, knockdown. The experiments were repeated three times.

***B***, HS578T, A549, and H1299 cells were treated with 30 μM cisplatin or vehicle for 24 h, then processed for IP of MDM2 and fluorescence IB. Fluorescence IP−IB detects stress-induced $PIP_2$ association with endogenous MDM2. The $PIP_2$ IB intensity was quantified, and the graph is shown as mean ± s.d. of $n = 3$ independent experiments.

***C***, PLA of MDM2-p53 in MDA-MB-231 cells treated with vehicle or 30 μM cisplatin for 24 h. The nuclear PLA foci of MDM2-p53 were quantified. $n = 30$ cells pooled from 3 independent experiments, 10 cells per experiment.

**Figure S2**

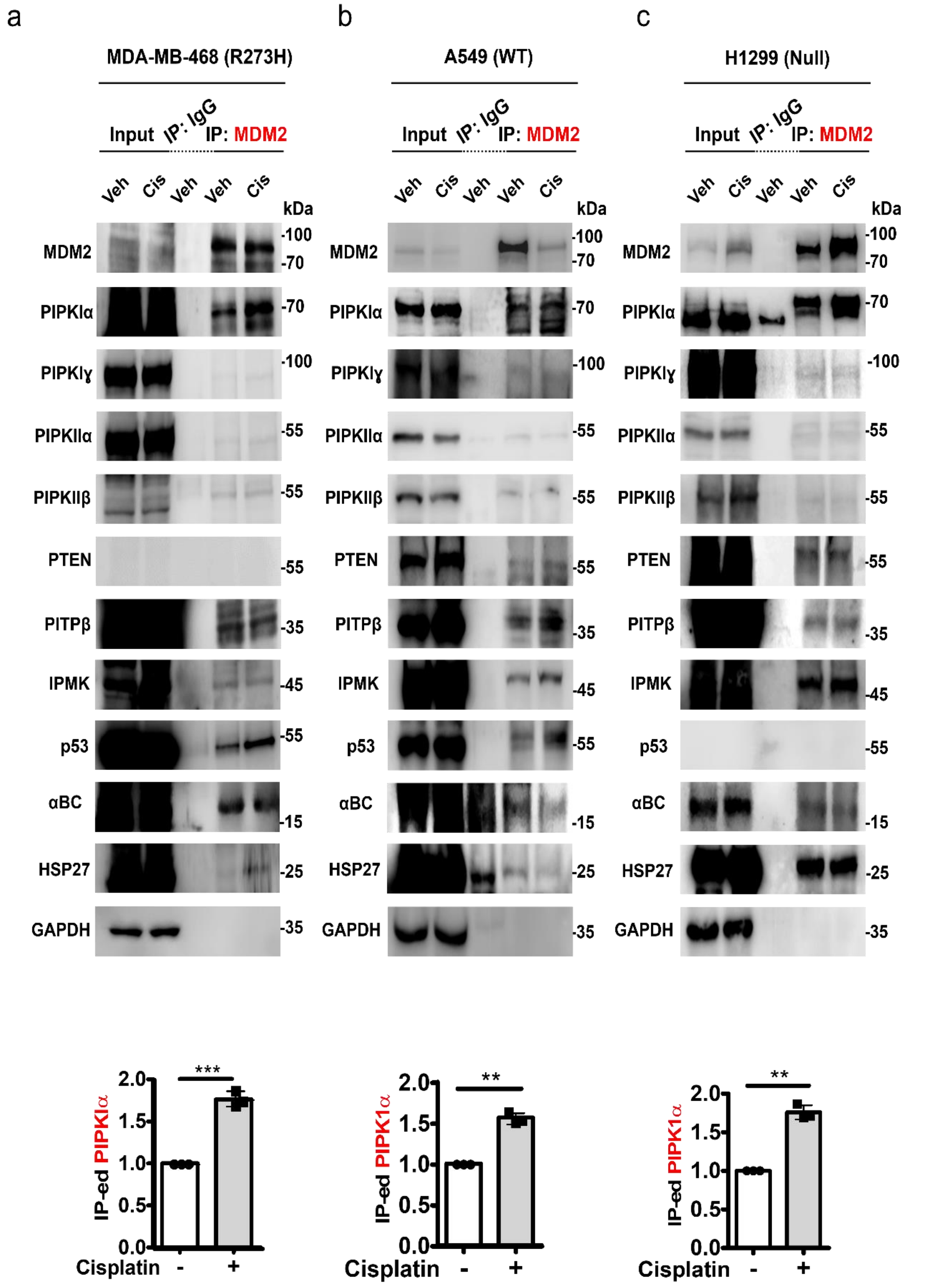

a
MDA-MB-468 (R273H)
b
A549 (WT)
c
H1299 (Null)
Input
IP: IgG
IP: MDM2
Veh
Cis
Veh
Veh
Cis
kDa
MDM2
PIPKIα
PIPKIɣ
PIPKIIα
PIPKIIβ
PTEN
PITPβ
IPMK
p53
αBC
HSP27
GAPDH
-100
-70
-55
-35
-45
-25
-15
IP-ed PIPKIα
IP-ed PIPK1α
Cisplatin
-
+
***
**
2.0
1.5
1.0
0.5
0.0

**Figure S2: MDM2 associates with PIP kinases and sHSPs in various cell lines**

***A-C*****,** Co-IP of endogenous MDM2 from MDA-MB-468 (*A*), A549 (*B*), and H1299 (*C*) cells treated with vehicle or 30 μM cisplatin for 24 h. The MDM2, PIPKIα, PIPKIγ, PIPKIIα, PIPKIIβ, PTEN, PITPβ, IPMK, p53, αBC, HSP27 and GAPDH co-IPed by MDM2 were analyzed by IB. Veh, vehicle; Cis, cisplatin-treated.

Figure S3

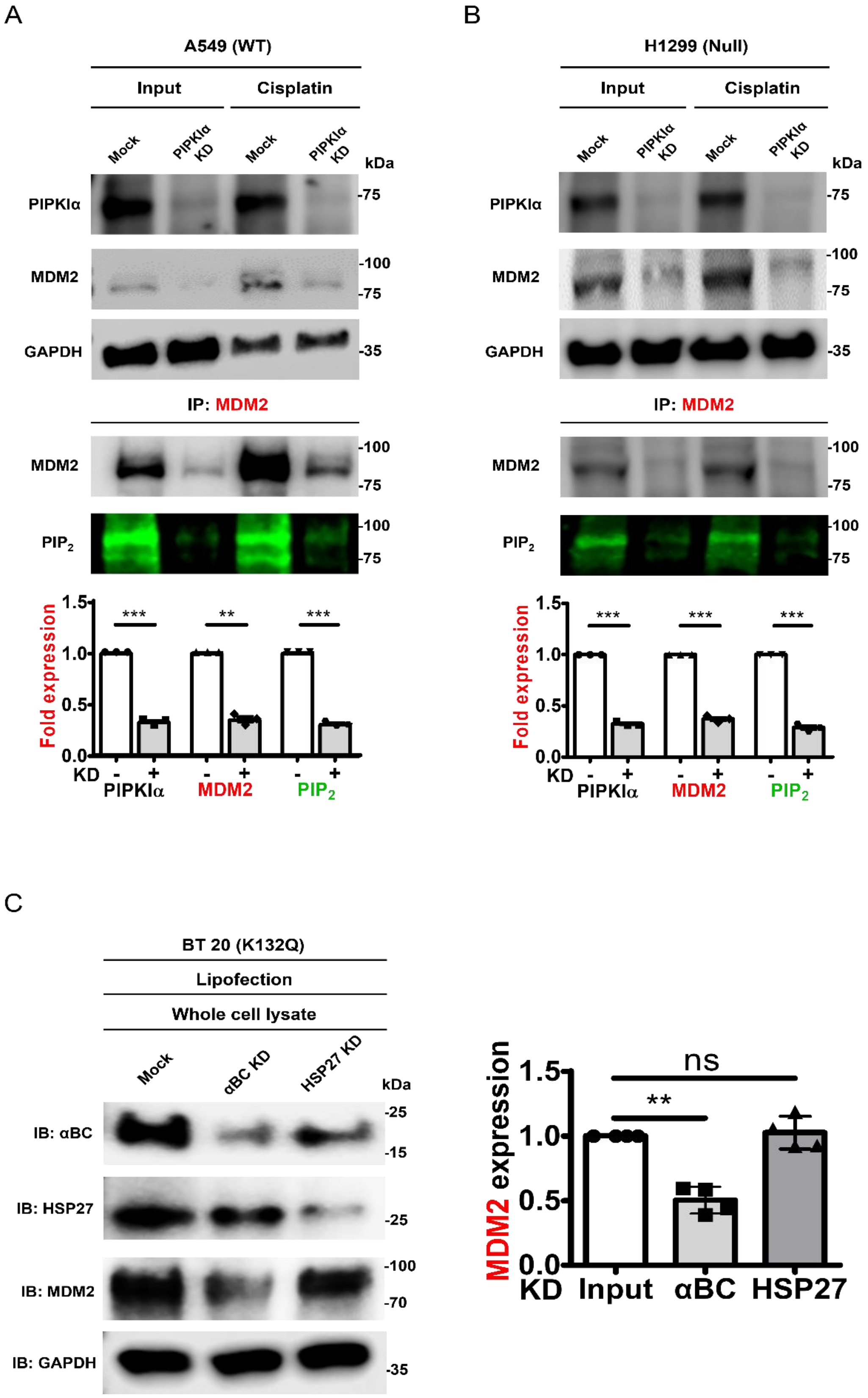
A
A549 (WT)
Input
Cisplatin
Mock
PIPKIα KD
Mock
PIPKIα KD
kDa
PIPKIα
-75
MDM2
-100
-75
GAPDH
-35
IP: MDM2
MDM2
-100
-75
PIP2
-100
-75
Fold expression
1.5
1.0
0.5
0.0
***
**
***
KD
-
+
PIPKIα
MDM2
PIP2
B
H1299 (Null)
Input
Cisplatin
Mock
PIPKIα KD
Mock
PIPKIα KD
kDa
PIPKIα
-75
MDM2
-100
-75
GAPDH
-35
IP: MDM2
MDM2
-100
-75
PIP2
-100
-75
Fold expression
1.5
1.0
0.5
0.0
***
***
***
KD
-
+
PIPKIα
MDM2
PIP2
C
BT 20 (K132Q)
Lipofection
Whole cell lysate
Mock
αBC KD
HSP27 KD
kDa
IB: αBC
-25
-15
IB: HSP27
-25
IB: MDM2
-100
-70
IB: GAPDH
-35
MDM2 expression
1.5
1.0
0.5
0.0
ns
**
KD
Input
αBC
HSP27

**Figure S3: PIPKIα and sHSP KD affect MDM2 stability**

***A*, *B*,** A549 (*A*), and H1299 cells (*B*) were transfected with siRNAs targeting PIPKIα. After 24 h, cells were treated with 30 μM cisplatin for an additional 24 h. IB analyzed the expression of the indicated proteins. The IBs of PIPKIα, MDM2, and $PIP_2$ were quantified. The graph shows mean ± s.d. of n = 3 independent experiments. KD, knockdown.

***C***, BT20 cells were transfected with siRNAs for αBC or HSP27. Expression of αBC, HSP27, MDM2, and GAPDH was analyzed by IB, and αBC and HSP27 IBs were quantified. The graph shows mean ± s.d. of n = 3 independent experiments. KD, knockdown. KD, knockdown.

**Figure S4**

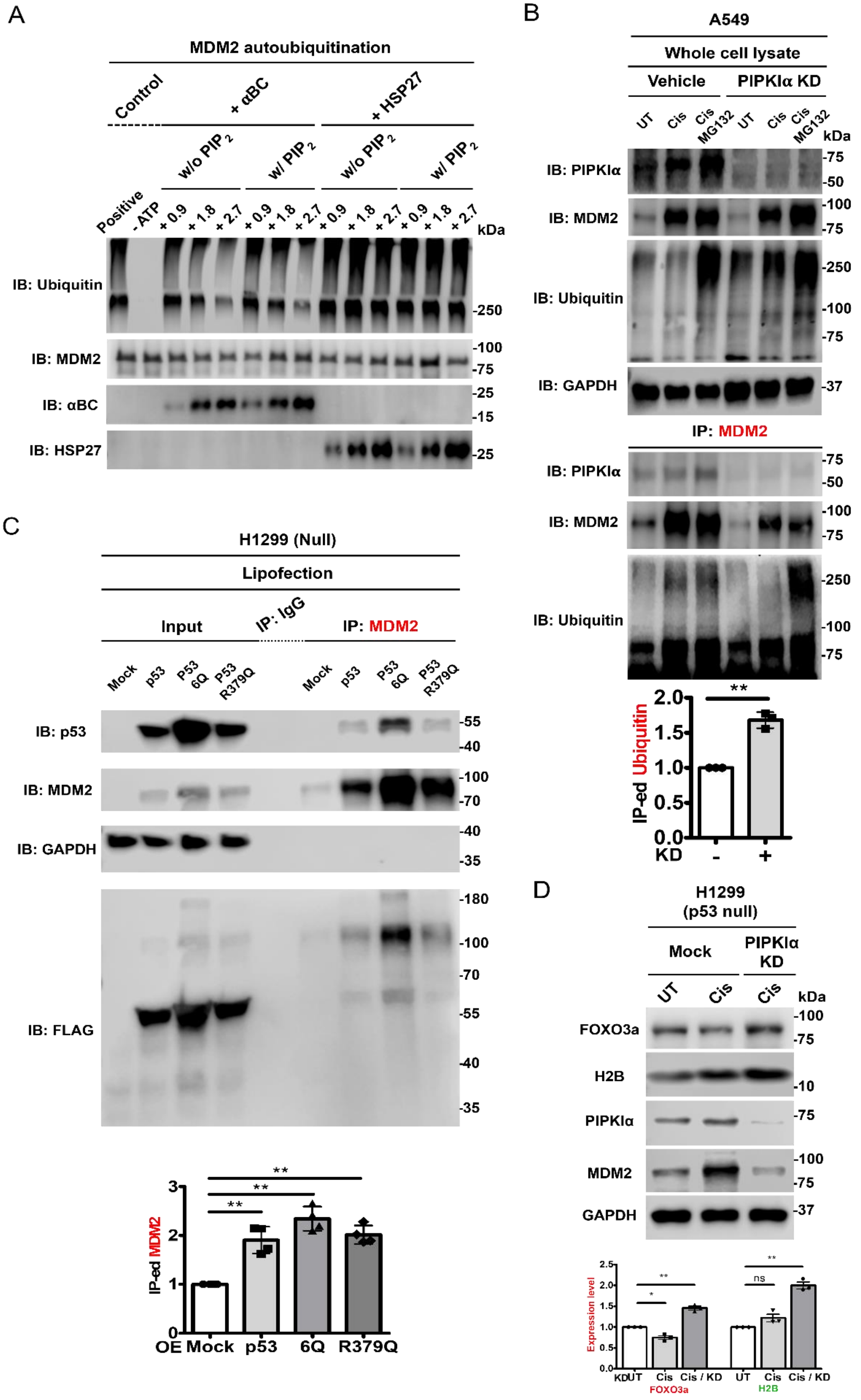
A
MDM2 autoubiquitination
Control
+ αBC
+ HSP27
w/o PIP 2
w/ PIP 2
w/o PIP 2
w/ PIP 2
Positive
- ATP
+ 0.9
+ 1.8
+ 2.7
kDa
IB: Ubiquitin
IB: MDM2
IB: αBC
IB: HSP27
-250
-100
-75
-25
-15
-25
B
A549
Whole cell lysate
Vehicle
PIPKIα KD
UT
Cis
Cis MG132
kDa
IB: PIPKIα
IB: MDM2
IB: Ubiquitin
IB: GAPDH
IP: MDM2
IB: PIPKIα
IB: MDM2
IB: Ubiquitin
-75
-50
-100
-75
-250
-100
-75
-37
IP-ed Ubiquitin
KD
-
+
**
2.0
1.5
1.0
0.5
0.0
C
H1299 (Null)
Lipofection
Input
IP: IgG
IP: MDM2
Mock
p53
P53 6Q
P53 R379Q
IB: p53
IB: MDM2
IB: GAPDH
IB: FLAG
-55
-40
-100
-70
-40
-35
-180
-100
-70
-55
-40
-35
IP-ed MDM2
OE Mock
p53
6Q
R379Q
**
**
**
3
2
1
0
D
H1299 (p53 null)
Mock
PIPKIα KD
UT
Cis
Cis
kDa
FOXO3a
H2B
PIPKIα
MDM2
GAPDH
-100
-75
-10
-75
-100
-75
-37
Expression level
KD
UT
Cis
Cis / KD
FOXO3a
H2B
*
**
ns
**
2.5
2.0
1.5
1.0
0.5
0.0

**Figure S4: Regulation of MDM2 ubiquitination and protein levels by αBC and PIPKIα**

***A***, For MDM2 autoubiquitination, 100 nM of E1 enzyme, 1 μM of E2 enzyme, 1 μM of MDM2, E3 ligase reaction buffer, 10mM of MgATP solution, and 100 μM of ubiquitin were incubated with different concentrations of αBC and HSP27 (0.9, 1.8, or 2.7 μM) in the absence or presence of $PIP_2$ (10 μM) for 1 h. IB analyzed Ubiquitin, MDM2, αBC, and HSP27. The experiments were repeated three times.

***B***, A549 cells were transfected with siRNAs for PIPKIα and treated with vehicle or 30 μM cisplatin for 24 h. After cells were treated with 10 μM of MG132 for 4 h, cells were harvested for IP of MDM2. IB analyzed the expression of the indicated proteins, and ubiquitin IBs were quantified. The graph shows mean ± s.d. of n = 3 independent experiments. KD, knockdown; UT, untreated; Cis, cisplatin-treated; Cis/MG132, Cisplatin/MG132 treated.

***C*, **H1299 cells were transfected with wild-type p53, p53 6Q and p53 R379 mutant for 48 h. Empty vector (Mock) was used as a negative control. Cells were subjected to Co-IP with MDM2. The expression of the indicated proteins, MDM2, p53, GAPDH, and FLAG tag, was analyzed and quantified by IB. The graph shows mean ± s.d. of n = 3 independent experiments.

***D***, H1299 cells were transfected with siRNAs for PIPKIα and treated with vehicle or 30 μM cisplatin for 24 h. After cells were harvested for IB. The expression of the indicated proteins was analyzed and quantified. The graph shows mean ± s.d. of n = 3 independent experiments. Mock, empty vector; KD, knockdown; UT, untreated; Cis, cisplatin-treated.
